# SciRide Finder: a citation-based paradigm in biomedical literature search

**DOI:** 10.1101/208959

**Authors:** Adam Volanakis, Krawczyk Konrad

## Abstract

There are more than 26 million peer-reviewed biomedical research items according to Medline/PubMed. This breadth of information is indicative of the progress in biomedical sciences on one hand, but an overload for scientists performing literature searches on the other. A major portion of scientific literature search is to find statements, numbers and protocols that can be cited to build an evidence-based narrative for a new manuscript. Because science builds on prior knowledge, such information has likely been written out and cited in an older manuscript. Thus, Cited Statements, pieces of text from scientific literature supported by citing other peer-reviewed publications, carry significant amount of condensed information on prior art. Based on this principle, we propose a literature search service, SciRide Finder (finder.sciride.org), which constrains the search corpus to such Cited Statements only. We demonstrate that Cited Statements can carry different information to this found in titles/abstracts and full text, giving access to alternative literature search results than traditional search engines. We further show how presenting search results as a list of Cited Statements allows researchers to easily find information to build an evidence-based narrative for their own manuscripts.

## 1. Introduction

More than 60,000 articles are deposited in PubMed each month, making literature search an increasingly difficult task^1^. A typical literature query consists of keyword based search by services such as Google Scholar, PubMed, Scopus or Web of Science^2–4^. The results typically consist of a list of titles and abstracts from documents that contain the query keywords. The scientist is then tasked with parsing through an extensive list of results, to extract information directly from titles/abstracts or to follow a link to the full document.

As such literature search can be burdensome, intelligent text mining of scientific publications has been seen as an alternative for extracting and organizing information from the ever-growing PubMed collection^5^. Sites such as iHOP or Chilibot mine field-specific knowledge by collating information regarding biomolecules from millions of PubMed publications^6,7^. Less field-specific services such as COLIL, provide a service showing comments in more recent research on older manuscripts^8^. These tools demonstrate that strategic text mining and intelligent filtering can lead to new, more efficient tools for biomedical literature search.

Strategic text mining can be used to separate relevant information from tangential text. For instance, because of legal restrictions, typical literature search engines operate on the remit of copyright-available titles and abstracts alone, whereas full text contains more pertinent information^9^. For instance, tools such as Biotext or Yale Image Finder allow searches in Figure or Table captions alone in order to identify relevant information only^10,11^. To understand what information is potentially irrelevant, it is necessary to identify portions of searchable documents that can be of more interest to the person performing literature search.

One major aim of literature search is to identify earlier papers to support the narrative presented in a new manuscript being written. Such narrative is constructed by citing findings, numbers, data and techniques from previous publications^12^. Such pieces of text are easily identifiable in scientific manuscripts since they are annotated with references to prior peer-reviewed publications which support the statement being made^12,13^. Therefore such statements in publications on previous literature, which we here call Cited Statements, offer succinct comments on prior art, whose information content is powerful enough to be used for article summarization^14^.

Here, we propose a simple strategy of improving text mining and literature search by creating a biomedical search corpus, which is constrained to such Cited Statements only. We show that Cited Statements can carry different information-retrieval data to these found in titles, abstracts and full text of documents they refer to, demonstrating that this methodology does not simply recapitulate information currently available in scientific search engines. Furthermore, we show how presenting results in the form of Cited Statement text, can offer easy access to information in several literature-search scenarios. We hope that our service, available at finder.sciride.org will offer a streamlined way for biomedical scientists to build evidence-based narratives for their own manuscripts.

## 2. Results/Applications

### 2.1 SciRide Finder as an alternative biomedical literature search platform

SciRide Finder offers an orthogonal literature search strategy to platforms such as PubMed or Google Scholar by focusing on Cited Statements only. In this manuscript, we refer to Cited Statement (CS) as any sentence from a peer reviewed publication containing citations to other manuscripts, which we refer to as Primary Research (PR) (Figure 1).

**Figure 1.**
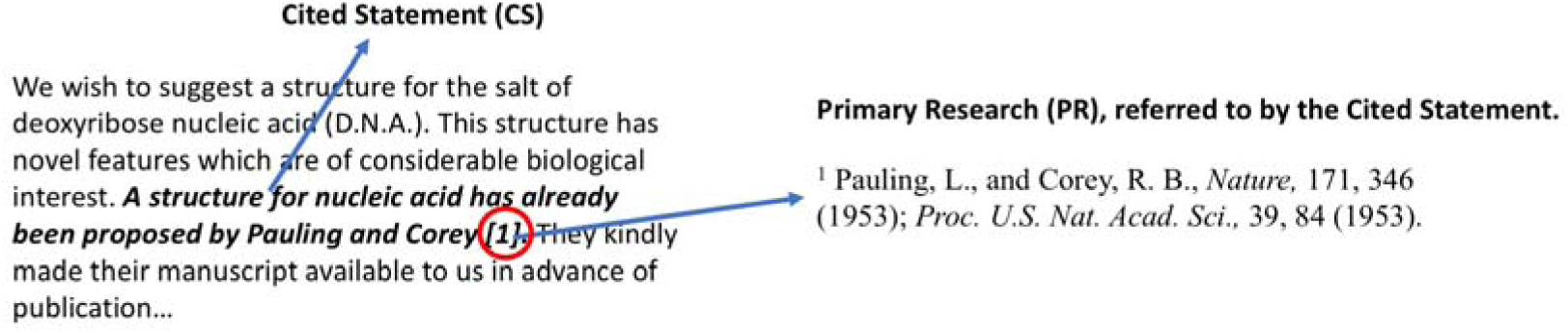
Example of a Cited Statement (CS) and Primary Research (PR). The CS is shown in bold in the excerpt on the left. PR which is referred to by the CS, is shown on the right. The text in the image was taken from the seminal paper by Watson and Crick in 1953, entitled ‘A Structure of Deoxyribose Nucleic Acid’.

We have extracted the CSs from all PubMed/Medline indexed documents where the copyright allowed for data mining and reproduction. To the best of our knowledge, the most suitable corpus for this task is the Open Access PubMed Central (OA PMC) dataset. It is a collection of open access journals from PubMed/Medline in standardized format. At the time of writing, there were approximately 1.7m publications in the OA PMC dataset, which is 6% of a total of more than 26m publications indexed in PubMed (or 15m if only citations with abstracts are to be considered^9^).

The OA PMC dataset downloaded via the NCBI ftp service forms the core of our dataset. Nevertheless, the ~1.7m OA PMC articles are only a subset of more than 4m web-formatted documents available via PMC^15^. There are more than 2m articles published after 1980 which are accessible via PMC ‘eyes-only’ subject to strict restrictions on machine access and heterogeneous publisher copyrights. We therefore extract such data manually if and only if the copyright situation is unambiguous.

We have set up a pipeline to collect data from the OA PMC and other publications in the public domain where copyright allows it (see Materials and Methods). At the time of writing, our data collection encompasses 1,786,322 peer-reviewed articles contributing 43,326,402 CSs. We make this corpus accessible via efficient Lucene-based search system as described in Materials & Methods. Here, we argue that our CS-based search system is a new literature search paradigm, distinct from traditional title/abstract and full text based methods.

The first major difference between traditional and CS-based search is the corpus employed to identify documents. In traditional systems, documents are retrieved if the query keywords are found within text of title, abstract or full text. On the other hand, searching by CSs identifies PR documents indirectly by text contained in other papers (Figure 2). CS offers an alternative commentary on the PR, by scientists who were generally not involved in the original study. To prove this point, we demonstrate that CSs can hold alternative information to titles/abstracts and full text of PR documents, described in section 2.2.

**Figure 2.**
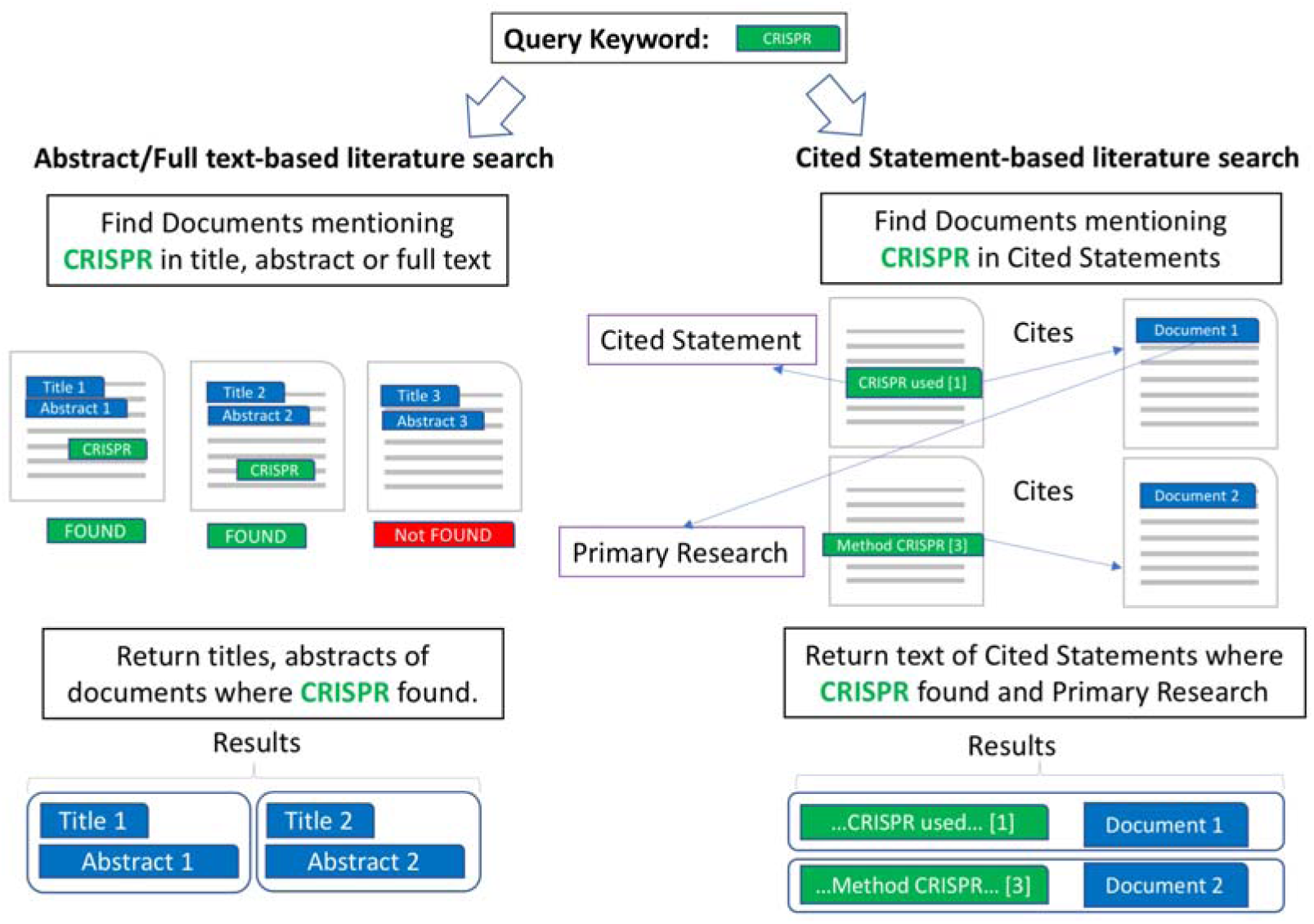
Contrasting the traditional literature search and Cited Statement-based literature search. Traditional literature search systems identify documents to be retrieved by keyword hits within them and present titles and abstracts as results (left). Cited Statement-based search identifies Primary Research documents by text from other publications and presents the citing text.

The second major difference between traditional and CS-based search is the presentation of results. In traditional literature search systems, results are presented as titles/abstract, more seldom as full text excerpts. In contrast, CS-based search returns the text which cites other documents. In this capacity, it identifies the information which was used to build the evidence-based narrative for a manuscript: scientific statements, numbers, data and techniques, all supported by prior publications. In many scenarios, these are the pieces of text scientists look for in the first place to build an evidence-based structure for their own manuscripts. To exemplify this, we present possible applications of presenting results as CSs in section 2.3.

### 2.2 Cited Statements can hold different information on documents to titles/abstracts and full text

We argue that CS-based search offers a novel way of retrieving documents, that can yield orthogonal results as compared to traditional search strategies. For this to be true, CSs must offer distinct information-retrieval data on the PR that would not normally be available by examining titles, abstract or even full text of PR document.

To quantify this, we identified 691,354 documents where we had CSs in our database referring to PR documents whose full text is available for text mining. For a given CS, we measured how many normalized words (stemmed, case-folded etc.) cannot be found in the title/abstract and full–text of the PR documents, which we refer to as ‘difference’. For each of 691,354 documents, we have identified the maximally different (as percentage) CS with respect to the title/abstract and full text of the PR document (Figure 3). These results demonstrate that for 83% of our 691,354 PR documents, there exists a CS which is at least 50% different to the title/abstract of the PR document. For 61% of PR documents, there exists a CS which is at least 25% different to the PR document full text. Therefore, for a significant proportion of publications, there exists a CS which offers information that would not be available through a title/abstract or full-text search on the PR document.

**Figure 3.**
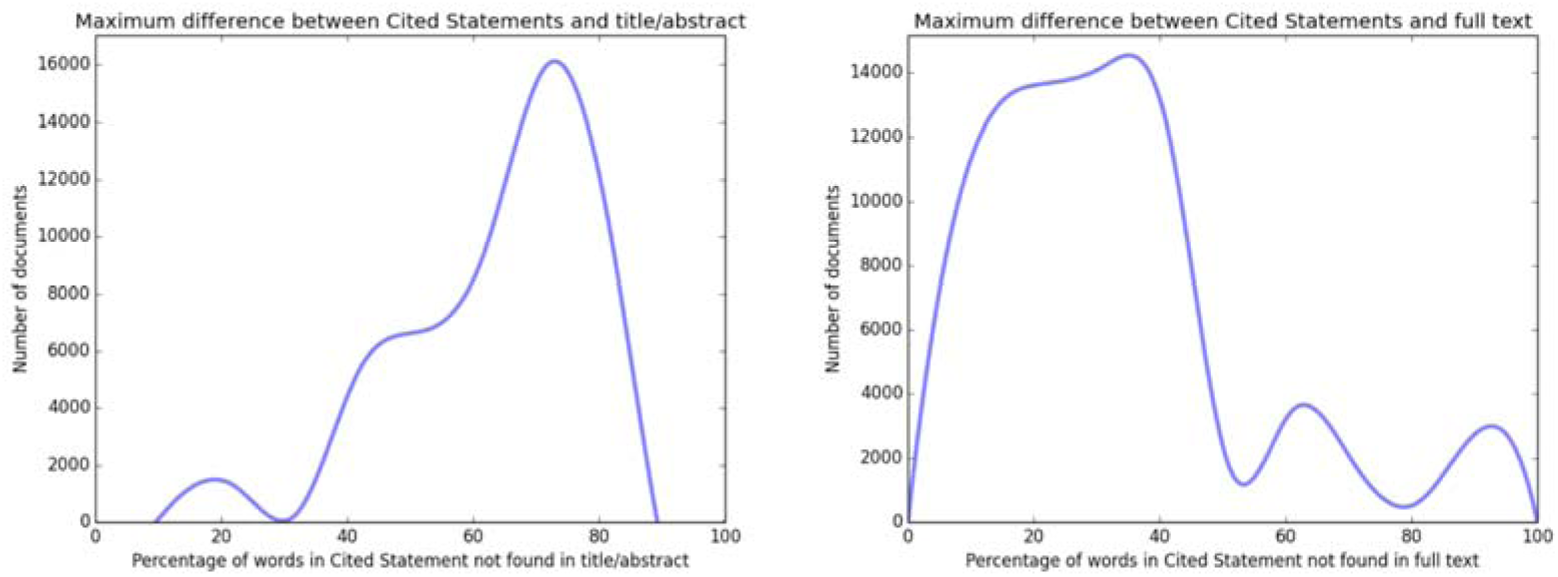
Maximum difference between CS and text of PR. For each of 691,354 documents, we identify the maximally different CS with respect to title/abstract (left) and full text (right) of PR. This stands to demonstrate that there are many PR documents, where there exists a significantly textually different CSs referring to them.

We have also found that CSs tend to be different from titles/abstracts not only in the extreme as above, but also on average (see Supplementary Figure 1). These results demonstrate that CSs can contribute different information on the PR manuscripts than title/abstract and full text. Therefore corpus constrained to CSs only can provide orthogonal results to those offered by search engines which identify documents directly by their titles/abstracts and full text.

### 2.3 Applications of the Cited Statement-based literature search

Search results from SciRide Finder do not consist of titles, abstracts or full-text excerpts as is typically the case in other services, such as PubMed or Google Scholar. Instead, in response to a query, we present a list of CSs, PR documents and papers where the information was found (Figure 4). To exemplify the utility of presenting results in this way we demonstrate possible literature search scenarios where CSs instantly provide information being sought after:

**Figure 4.**
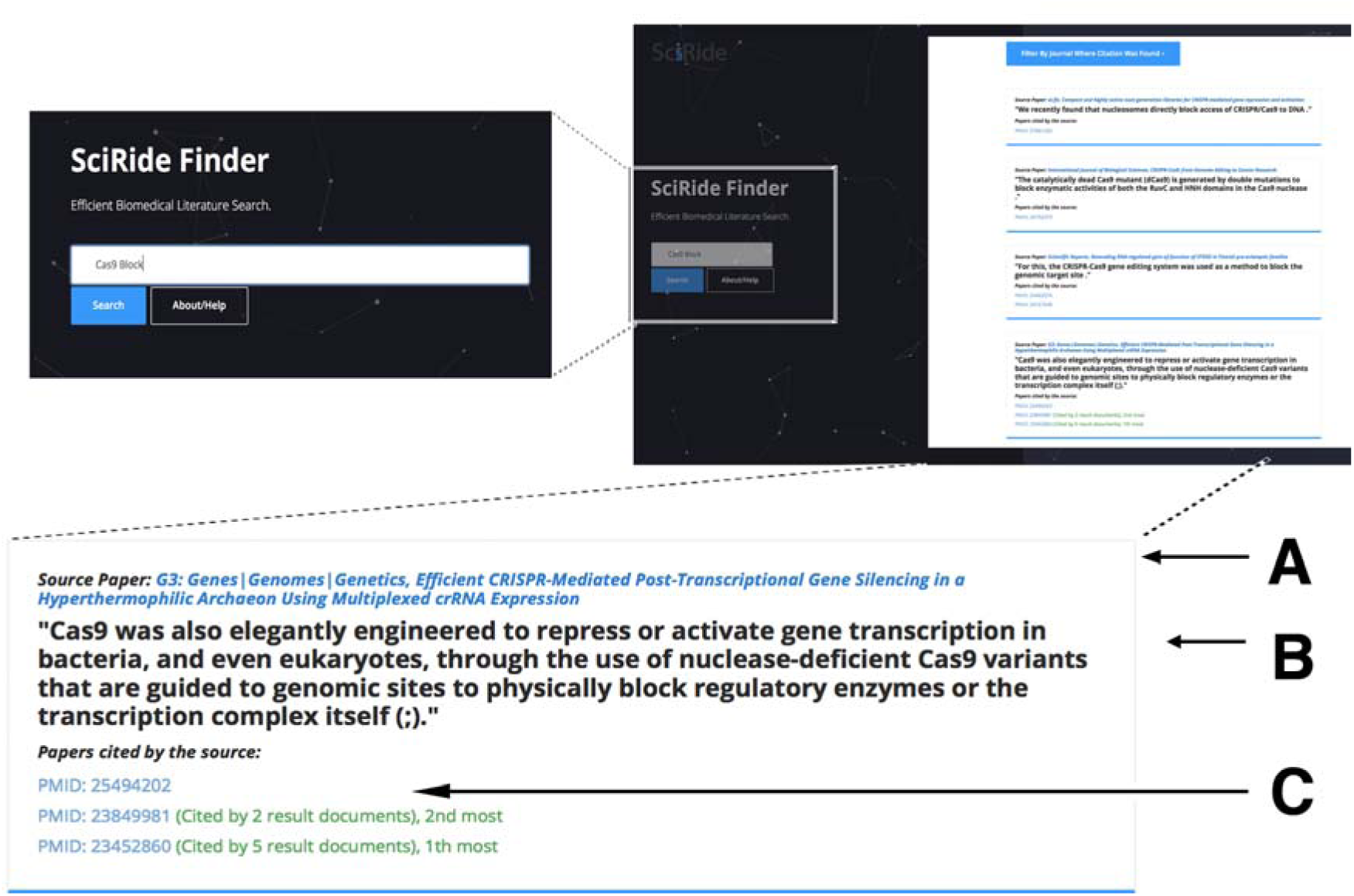
SciRide Finder search example for “Cas9 block” statements. Each result consists of the CS (**B**), the title of the paper that the statement appears in (**A**) and the PR sources that the statement is based on (**C**).

#### Identifying citations supporting general knowledge

It is often problematic to identify citations supporting a well-known fact. For instance, the controversy surrounding the link between vaccines and autism is well known, but identifying studies discrediting it is not trivial. Searching SciRide Finder for “autism” “vaccine” and “discredited” would return results that debunk the notorious 1998 publication in Lancet by Wakefield and colleagues.

#### Identifying datasets

Datasets used in publications are rarely cited in titles and abstracts, rather being hidden in Methods sections. As an example, searching SciRide Finder for the terms “ChIP-seq” “HeLa” and “Pol II” would return the publications that have used datasets of RNA Polymerase II (Pol II) Chromatin Immunoprecipitation sequencing (ChIP-seq) experiments in HeLa cells but also the original source of these datasets (Supplementary Figure 2), thus facilitating the retrieval of the datasets.

#### Research technique identification

Similarly to datasets, specific techniques used are rarely available in abstracts and titles. Nevertheless, identifying publications which employ a given technique or software is indispensable for reuse of protocols. For example, the newly developed CRISPR/Cas9 genome editing method has an alternative usage as a block for gene transcription. A PubMed search for the terms “Cas9” and “Block” returns 47 publications (at the time of writing) and there is no way of knowing how and in what context this method was used in each paper without reading all manuscripts. The same search in SciRide Finder (Figure 4) provides a list of publications where this technique was used in context. This allows us to identify publications describing the method, the theory behind it, protocols used, and the original research.

#### Mapping connections betwen keywords and publications

SciRide Finder allows for searching for two or more terms appearing together, their context and the original research. For example, a CS-search for ‘mRNA export’ and ‘transcription’ would identify only the statements in which the two keywords appear together (Supplementary Figure 3). Mapping such connections between keywords and citations can be of particular interest for creating knowledge maps by the text mining community.

## 3. Conclusion

We have created a biomedical search service based on information content from CSs. These short pieces of text build the evidence-based narrative for a given manuscript and provide a reflection of knowledge contributed by previous publications. We have shown that constraining the search corpus to CS only, can be a viable alternative to conventional search methodologies as it provides different information from titles/abstracts and full text. Furthermore, presenting results as CSs, is beneficial in many areas of scientific literature search, whose major part is aimed at identifying evidence-based pieces of text to be used in future publications.

Previous search methodologies, such as Google Scholar, aim to index all the information available on documents even if the publication itself is not in the public domain. On the contrary, our service indexes only a very well-defined subset of the full-text articles, namely the CSs. We currently extracted ~43m CSs which contain comments on 34% (or 57%, if publications without abstracts are to be omitted^9^) of all of PubMed articles. This proportion should only increase as more publications become open access and repositories become legally and technically unified for systematic text mining^15^. Furthermore, since results of our service come solely from open access publications, it would follow that such manuscripts would be more readily cited as their content is freely accessible for scientists and search engines alike^16^.

In summary, our system introduces on open-access, CS-only paradigm in literature search. Current manifestation of this paradigm, SciRide Finder, offers an orthogonal approach to reduce the burden currently associated with specific information retrieval in biomedical literature. We hope that our service will facilitate the efforts of researchers looking for Cited Statements, to build an evidence-based narrative for their own publications.

## 4. Materials & Methods

### 4.1 Data Collection for the base system – PMC Open Access Dataset

The OA PMC corpus was downloaded from the NCBI FTP website (ftp.ncbi.nlm.nih.gov) and divided into sentences using Natural Language Toolkit (nltk.org) and a custom set of heuristics, such as splitting text on terminal period ‘.’, removing the ‘.’ from short-hands such as ‘et al.’, ‘ca.’ and normalizing the scientific names (‘H. Pylori’). We identified the sentences containing citations as these having the <xref> tag with attributes pointing to references section (as opposed to non-bibliographic elements such as Tables and Figures). Rules were created for special cases where the citation pertaining to a sentence occurs after its terminal period. Each CS derived in this way contains the citing sentence, identifiers of cited articles (DOI or Pubmed ID) and the metadata on the manuscript it was derived from (journal title and article title). The system was set up to perform updates of this base dataset on a monthly basis.

### 4.2 Data Collection Beyond the PMC Open Access Dataset

We augment the information from the base-dataset manually from the ‘eyes-only’ documents where there was an unambiguous copyright situation on reproducing pieces of work in a normal citation scenario. Furthermore, it is sometimes possible to find the author-submitted PDF version of the document. These are documents available via platforms such as BioArxiv or author homepages. Whenever we could not identify a PMC version of an article, we attempted a PDF doi search. When a PDF document was found in such a way, we extracted information from it using PDFExtract tool from CiteSeer. PDFExtract is a utility which is capable of extracting portions of a PDF-formatted scientific document and present them in machine-readable plain text. Since information presented in such format is still very heterogeneous, we had to create different sets of rules to interpret the PDF-extracted plain text, which mostly involved detecting if the citations are number-based or author-name based.

### 4.3 Text Retrieval System

The Cited Statements are stored for rapid extraction in a Lucene-based system which was previously shown to be a robust search engine for biomedical applications^17^. Since scientific documents are by and large written in English^18^ we have employed standard English analyser and stemmer as parameters of retrieval. We only perform searches on the text of the CS record, disregarding metadata of the full article it was retrieved from.

Documents are retrieved given a set of keywords to match the text of the CS. A post-processing step after document retrieval is introduced, where we count the number of shared citations between resulting documents. The documents are sorted in descending order firstly by the relevance score of the Lucene system (normalized to one decimal point) and secondly by the number of shared citations. This assures that statements on highly cited papers which are similar within the normalized value of the text-relevance score, are displayed first. The literature search service is available as a web service at http://finder.sciride.org. The text-mining of the CS corpus is available through an API which is described on the website.

### 4.4 Information content comparison

We measured how different the CSs are to PR document titles, abstracts and citations. Since a typical search engine operates on the remit of keywords, we have created a textual fingerprint for each CS, title/abstract and full text. Each fingerprint was a set of case-folded, stemmed and stop-word-free normalized words without duplicates.

For each PR document, we have collected three elements: its title/abstract, full text and a list of CSs referring to it. We have created a textual fingerprint for each title/abstract, full text and each CS, which was supposed to emulate a typical corpus employed by an information retrieval system.

To produce a fingerprint for a given piece of text, we split it into word tokens using the NLTK toolkit. We case-folded each word, and removed any punctuation (keeping special symbols such as Greek letters). We removed all stop-words (as defined by the NLTK corpus). Finally each word was stemmed so as to minimize mismatches in subtle inflection forms^19^. We did not keep word duplicates, thus for each text element (such as title/abstract), this resulted in a non-redundant list of normalized words.

A typical information retrieval algorithm can be expected to perform such text-normalization operations on a given document. Thus, it is reasonable to assume that if text-normalized fingerprints share many words, the information retrieval algorithm would treat them as contributing similar information and yield similar results. Therefore, the number of different normalized words between CS and PR title/abstract and full text was taken as a measure if CS contribute new information on PR.

Comparing two fingerprints (e.g. CS versus full-text) consisted of counting how many text-normalized words are found in one fingerprint but not the other.

## Author Contributions

AV and KK designed the experiments wrote the manuscript and prepared the figures. KK wrote the text mining algorithms and created the finder.sciride.org website.

## Additional information

The authors declare no competing financial interests related to this work and any material used in this study.

